# Overnight caloric restriction prior to cardiac arrest and resuscitation leads to improved survival and neurological outcome in a rodent model

**DOI:** 10.1101/786871

**Authors:** Matine Azadian, Guilian Tian, Afsheen Bazrafkan, Niki Maki, Masih Rafi, Monica Desai, Ieeshiah Otarola, Shuhab Zaher, Ashar Khan, Yusuf Suri, Minwei Wang, Oswald Steward, Yama Akbari

## Abstract

While interest toward caloric restriction (CR) in various models of brain injury has increased in recent decades, studies have predominantly focused on the benefits of chronic or intermittent CR. The effects of ultra-short, including overnight, CR on acute ischemic brain injury, however, are not well studied. Here, we show that overnight caloric restriction (75% over 14 h) prior to asphyxial cardiac arrest and resuscitation (CA) improves survival and neurological recovery, including complete prevention of neurodegeneration in multiple regions of the brain. We also show that overnight CR normalizes stress-induced hyperglycemia, while significantly decreasing insulin and glucagon production and increasing corticosterone and ketone body production. The benefits seen with ultra-short CR appear independent of Sirtuin 1 (SIRT-1) and brain-derived neurotrophic factor (BDNF) expression, which have been strongly linked to neuroprotective benefits seen in chronic CR. Further exploration into the precise mechanisms involved in ultra-short CR may have significant translational implications. These findings are also of importance to basic sciences research as we demonstrate that minor, often-overlooked alterations to pre-experimental dietary procedures can significantly affect results, and by extension, research homogeneity and reproducibility, especially in acute ischemic brain injury models.

## Introduction

For almost a century, caloric restriction (CR) has been shown to have a multitude of health benefits in both humans and animals^1^. CR is defined as a reduction in caloric intake and can be daily, life-long, or intermittent. Many of the benefits appear to target aging, which includes prolongation of lifespan^2,3^ and improvements in age-related deficits of learning^4^ and memory^5^. In addition to its effects on aging, CR has been shown to be beneficial in various models of neurological diseases, most notably Alzheimer’s^6^, Parkinson’s^7^, and epilepsy^8^. Recently, CR has also been shown to have neuroprotective effects in various models of traumatic brain injury^9,10^ and stroke^11^.

Several studies on CR have proposed to explain its cellular and molecular mechanisms of action on the brain, which involve a wide range of pathways, including metabolic, inflammatory, oxidative stress, and cellular regenerative mechanisms^12,13^. These studies, however, have predominantly focused on long-term CR, which is clinically impractical for a variety of reasons, including issues of adherence and impracticality towards acute brain injuries. As a result, more interest has been garnered toward short-term CR, which can range from days to months and has been shown to have both cardio- and neuroprotective benefits^12,14^. In one study, CR for 3 days significantly reduced infarct volume after severe focal stroke in a rodent model^15^. In others, CR for 14 days induced brain ischemic tolerance in a rodent model of middle cerebral artery occlusion via upregulation of SIRT-1^16^; and intermittent CR every 24 h for 3 months upregulated BDNF in rodents, which protected neurons against excitotoxic injury^17^. Although long-term CR seems to involve a multiplicity of cellular pathways, it appears that short-term CR has thus far been linked to SIRT-1 and BDNF pathways, as several additional studies have pinpointed these downstream mechanisms, particularly in models of traumatic brain injury and focal stroke^17–19^.

Studies involving ultra short-term caloric restriction of a single day or less are sparse. We were interested in the effects of a simple overnight caloric restriction prior to global ischemic insult, which has not been previously investigated. Pre-clinical analysis of such phenomena may provide simple and translatable approaches to potentially ameliorate recovery following focal ischemia, which affects over 795,000 people per year in the United States^20^, and global ischemia, which affects over 550,00 people annually in the United States^21^. In this experiment, we induced global ischemia by CA in rats that were calorically restricted (75%) overnight for 14 h and assessed for changes in outcome. In attempt to better understand such changes, we measured levels of glucose, insulin, glucagon, corticosterone, and ketone bodies in the blood, in addition to SIRT-1 and BDNF expression in the brain.

## Materials and Methods

### Animals

Adult male Wistar rats (Charles River Laboratories, Wilmington, MA) weighing 300-370g were used in this study. The animals were housed under standard conditions (23±2°C, 60–70% relative humidity, 12 h light and dark cycles; free access to food and water). Animals typically arrived 2 weeks prior to experiments and were handled daily for 5 min to promote habituation and reduction of stress levels. All animal procedures were approved by the University of California Animal Care Committee (Irvine, CA) and conformed to the recommendations of the American Veterinary Medical Association Panel on Euthanasia.

### Dietary restriction

Rats were divided into two groups, control (n= 14) and calorically restricted (CR; n=14). The control group had unlimited access to food during the entire experiment. Rats from the CR group were fed 25% of the average daily food intake of the control group. Average daily food intake was calculated as weight of standard laboratory chow pellets consumed per day per rat. CR rats were calorically restricted overnight, starting at 6:00pm, approximately 14-hrs prior to surgical procedures at 8:00am on the following morning and 18-hrs prior to cardiac arrest at 12:00pm. Capillary blood ketone (β-hydroxybutyrate) levels were measured the morning after caloric restriction prior to surgical procedures with a Precision Xtra® System (Abbott, Princeton, NJ). Rats in the CR group were allowed to resume *ad libitum* feeding during the recovery period following surgical and cardiac arrest procedures. Both groups had *ad libitum* access to water throughout the experiment.

### Cardiac arrest experiment

On the day of CA, rats were endotracheally intubated, connected to a TOPO^TM^ mechanical ventilator (Kent Scientific, Torrington, CT), and maintained under 2% isoflurane anesthesia carried by 50% O_2_ and 50% N_2_ gas during the surgical preparations leading up to CA. The femoral artery and vein were cannulated to monitor blood pressure and heart rate and to allow for the intravenous (i.v.) administration of medications. While under mechanical ventilation, positive end expiratory pressure was maintained at 3 cmH_2_O and body temperature was monitored with a rectal probe and maintained at 37°C. Cardiac arrest was induced via an 8-minute duration of controlled asphyxia followed by cardiopulmonary resuscitation (CPR) until return of spontaneous circulation (ROSC) as previously described^22^. No isoflurane anesthesia was administered during or after CPR for the remainder of the experiment. Approximately 250 μL of arterial blood was collected 10 minutes before asphyxia and 10 minutes after ROSC. Blood gas levels, in addition to blood glucose, was measured at both timepoints with an i-STAT® System (Abbott, Princeton, NJ). Vessels were decannulated, and when spontaneous respirations were adequate, rats were extubated. Physiological parameters including blood pH, pO2 and pCO2 were monitored during surgery. There was no significant difference between CR and control groups either before or after CA (supplemental figures).

### Post-cardiac arrest care

Normal saline (5mL) and Ringer’s lactate (5mL) was administered subcutaneously (s.c.) 5 hours after CPR to limit dehydration until the rats resumed water consumption independently. Prophylactic cefazolin (45 mg/kg) was administered to limit any risk of infection. One-fourth cup of HydroGel (ClearH_2_O, Portland, ME) and 10 pellets of standard laboratory chow was soaked in water and placed near the rats’ mouth and throughout the cage until they recovered and resumed independent consumption of standard chow. At 24h post-CA, capillary blood ketone levels were again measured. Rats were re-examined every 24h thereafter to ensure proper hydration and food consumption. Both CR and the control group underwent analogous surgical, cardiac arrest, and post-cardiac arrest procedures. All rats were permitted *ad libitum* access to food and water following cardiac arrest.

### Neurological evaluation

Neurological recovery was quantified using the Neurological Deficit Scale (NDS; Table 1). The NDS consists of components that measure arousal, brainstem function, motor and sensory activities, as described previously^22^. Neurological recovery was evaluated at 4, 24, 48 and 72h post-ROSC by well-trained personnel who were blind to dietary groupings.

**TABLE 1.**
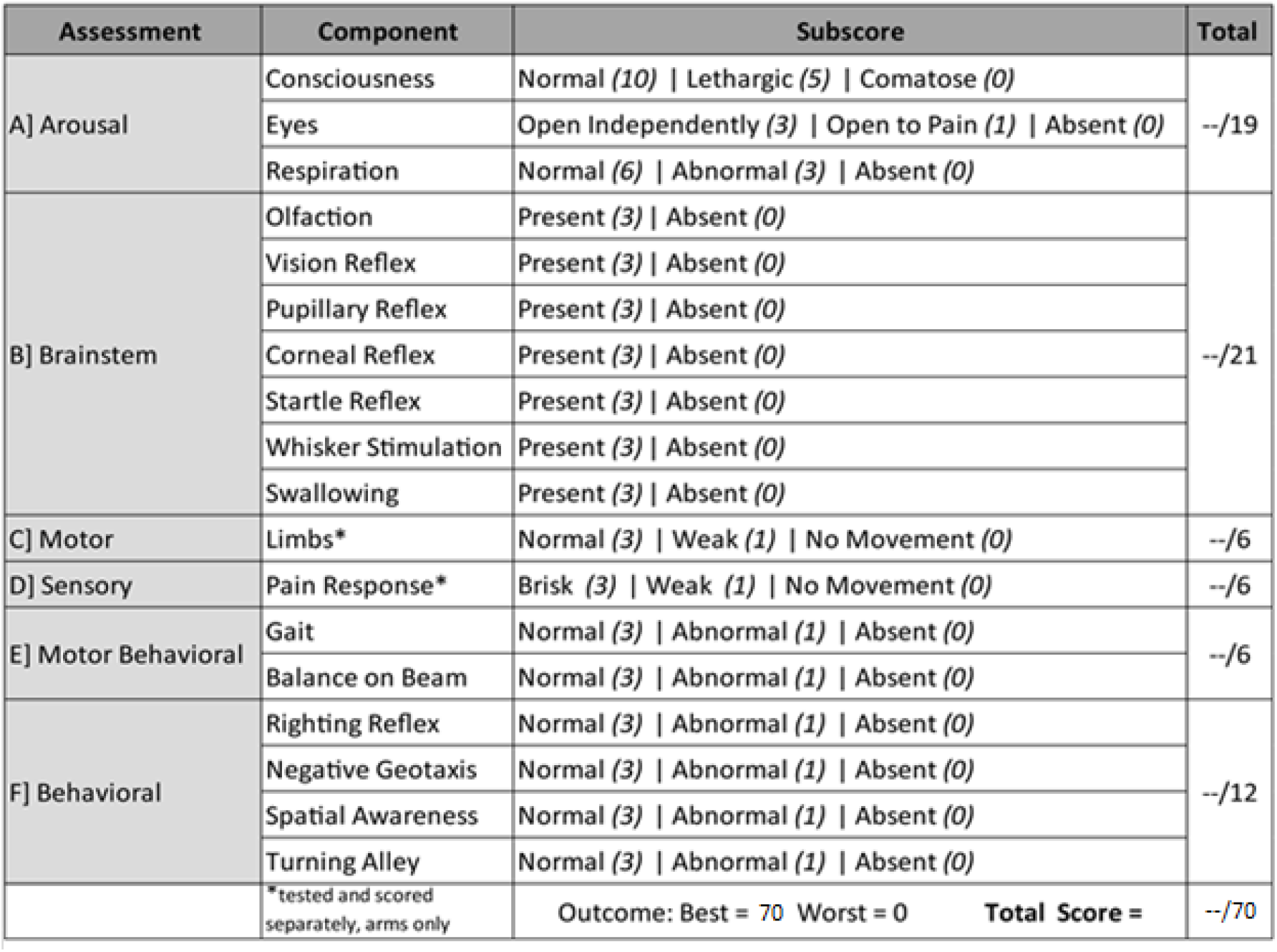
Neurological Deficit Scale (NDS)

### Brain tissue collection

At 72h following CA, survived rats were anesthetized with sodium pentobarbital and perfused transcardially with 0.9% NaCl solution followed by 0.1 M phosphate buffered saline, pH 7.4. Brains were separated at the mid-sagittal plane into left and right hemispheres. The left hemisphere was post-fixed in 4% PFA for 24 hrs at 4°C and cryoprotected in 30% sucrose for 4 days. It was then frozen in optimal cutting temperature (OCT) embedding medium and stored in −80°C until sectioned. The right hemisphere was flash frozen in dry ice and stored in −80°C until homogenization for western blot analysis.

### Histologic analysis

Left brain hemispheres frozen in OCT were coronally sectioned at 30 *μ*m using a cryostat (Microtome HM 505N). Sections were stored in serial order in a 96-well plate in 1 x PBS with sodium azide at 4°C. Fluorojade-B staining was used to scan and identify neuronal degeneration at 72h post-CA. To conform to the stereological standards of systematic random sampling, sections from figure 10 (bregma, 3.24 mm) to figure 155 (bregma, −14.64 mm) in the rat brain atlas (*The Rat Brain in Stereotaxic Coordinates, 6^th^ edition*) were chosen at a 330 μm interval (one of every twelve sections) covering a total 17880 μm scanned area of potential neurodegeneration. Selected sections were stained for 20 minutes in 0.0004% Fluorojade B solution (EMD Millipore, Billerica, MA, USA) after 10 minutes incubation with 0.06% potassium permanganate. Sections were screened under the microscope and those with Fluorojade-B positive neurons were marked for cell counting. To further avoid biased sampling, two additional sections were chosen at – 90 *μ*m anterior and +90 *μ*m posterior of the original marked section.

### Cell counting

Images of sections were obtained on a Nikon Eclipse Ti-E microscope (Nikon Corporation, Tokyo, Japan) under standardized conditions, including exposure time, gain, and resolution. Square sampling fields were numerated and placed on images to encompass areas with Fluorojade-B positive neurons. A random integer generator was utilized to select half of the numerated sampling fields for cell counting. Unbiased counting frames, modified from West^23^, were used to manually count cells, in which Fluorojade-B positive neurons that were partially or entirely within the top and right borders and did not intercept the bottom or left borders were considered to be in the counting frame and counted. This method ensured that Fluorojade-B positive neurons, regardless of size, shape, and orientation were not counted more than once. All images were blindly analyzed by three trained personnel using ImageJ Plugin “Cell Counter” (NIH, Bethesda, MD).

### Measurement of blood serum analytes

Arterial blood samples collected during the CA experiment were processed in EDTA-coated tubes with 25 μL aprotinin. After centrifugation (1,000 x g, 15 min), serum samples were aliquoted and stored at −80°C until use for measurement of analytes.

Serum concentrations of corticosterone, glucagon, and insulin were simultaneously determined by using a magnetic bead assay (Milliplex MAP Rat Stress Hormone/Metabolic Panel, Millipore, Billerica, MA, USA). All procedures were performed according to manufacturer’s instructions, at room temperature and protected from light. Samples were analyzed in a Luminex MAGPIX system (Millipore Sigma, Burlington, MA). Analyte concentrations were calculated using Analyst software (Millipore Sigma, Burlington, MA) with a five-parameter logistic curve-fitting method.

### Western blot analysis

Brain segments in the right hemisphere (cerebellum for SIRT-1 analysis; hippocampus for BDNF analysis) were sonicated in PBS containing Pierce Protease Inhibitor (Cat #88665, Fisher Scientific, Hampton, NH), assayed for total protein concentration, and then mixed with SDS sample buffer. The resulting samples were resolved by SDS-PAGE (8% polyacrylamide for SIRT-1 analysis; 16% polyacrylamide for BDNF analysis) and transferred onto PVDF membranes. The antibodies used were: ECL ™ anti-rabbit IgG (Cat #NA 934-1ml, GE Healthcare, Chicago, IL; 1:5000 dilution), mouse anti-beta tubulin (Cat# E7, Developmental Studies Hybridoma Bank of University of Iowa, Iowa City, IA), IRDye 800cw donkey anti-rabbit (Cat #32212, Li-Cor Biosciences, Lincoln, NE; 1:10,000 dilution). For BDNF analysis the primary antibody used was rabbit anti-BDNF N-20 (Cat #SC-546, Santa Cruz Biotechnology, Dallas, TX; 1:1000 dilution). For SIRT-1 analysis the primary antibody used was rabbit SIRT-1 (Cat # 07-131, Millipore Sigma, Burlington, MA; 1:500 dilution). The immunoreactive bands were detected using Amersham™ ECL Select™ detection reagent (Fisher Scientific, Hampton, NH) according to the manufacturer’s instructions. Bands were analyzed with ImageJ (NIH, Bethesda, MD).

### Statistical analysis

Data analysis was performed using IBM SPSS Statistics Software (V21; IBM Corporation, Armonk, NY) and GraphPad Prism (V6.0; GraphPad Software Inc., La Jolla, CA). Specific statistical tests utilized are noted accordingly below. Appropriate *post-hoc* tests were used as described. Data are presented as mean ± standard deviation unless otherwise noted. *, *p* < 0.05; **, *p* < 0.01, were considered significant.

## Results

### CR induces normoglycemia and inhibits stress-induced hyperglycemia

To evaluate the effect of 14-hrs of CR on glycemia, a contributable outcome factor in countless models of brain injury, we measured arterial blood glucose levels 10 min prior to and after CA. As expected, surgical preparation for the CA experiment resulted in control *(ad lib)* rats exhibiting stress-induced hyperglycemia, with glucose levels above normal. However, as shown in Figure 1, the CR group exhibited significantly lower blood glucose levels prior to CA (mean ± standard deviation [SD], 142.3 ± 12.8 mg/dL; n = 14) in comparison to control (219.5 ± 16.4 mg/dL; n = 12; *p*<0.01). Blood glucose levels remained stable after cardiac arrest in the CR group (141.8 ± 42.8 mg/dL) but not in the control group (277.7 ± 30.6 mg/dL), which exhibited a significant increase in glucose levels compared to pre-CA control values (*p*<0.01). These results suggest that rats undergoing asphyxial CA display a stress-induced hyperglycemic response that is inhibited in entirety by 14-hrs of caloric restriction.

**FIG. 1.**
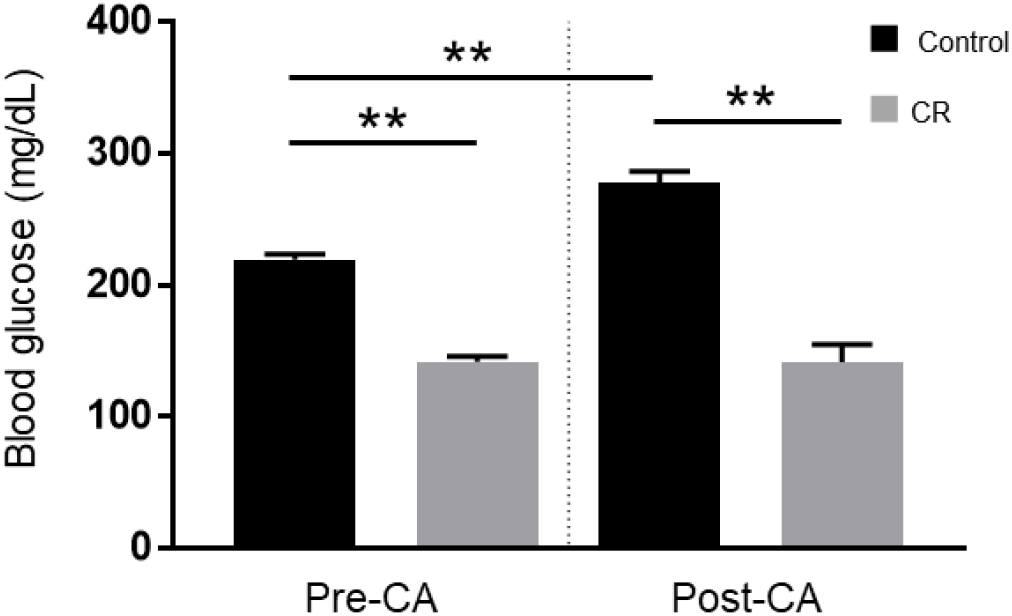
Arterial blood glucose of control vs. CR rats. Blood glucose was measured 10 minutes prior to CA and 10 minutes after resuscitation. While control (*ad lib*) rats demonstrate stress-induced hyperglycemia, CR significantly lowers and stabilizes blood glucose through the period of CA + CPR. ** *P* < 0.01; by oneway analysis of variance. CA, cardiac arrest; CON, control; SEM, standard error of the mean.

### CR improves survival after cardiac arrest

Given the overall severity of our CA model attributable to an 8-min duration of asphyxia, we expected several mortalities. In the control group, two rats failed to achieve ROSC during CPR and an additional two deceased at 24 h and 48 h post-ROSC. Remarkably however, all rats in the CR group successfully resuscitated and survived the full term of experimentation (72 h). To assess whether such an acute duration of caloric restriction affects survivability, a Kaplan-Meier survival analysis (Kaplan & Meier, 1958) was conducted. There was a statistically significant difference in survival distributions for the control versus CR group (*p*<0.05; Fig 2). This suggests that 14-hrs of caloric restriction may have significant downstream cardioprotective and neuroprotective effects during global ischemic injury.

**FIG. 2.**
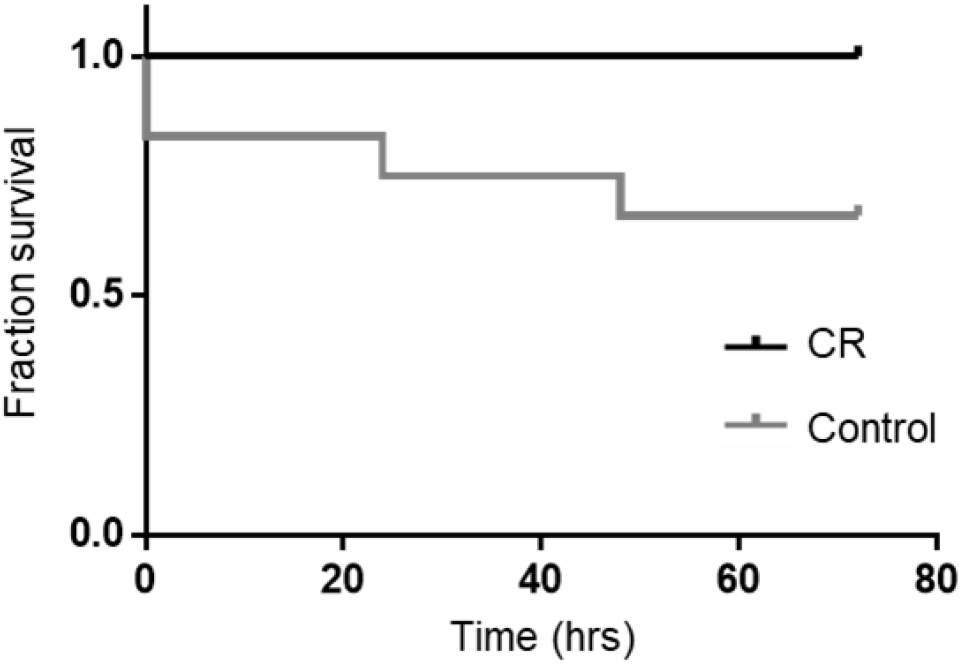
Fraction survival of post-CA rats. No mortalities occurred in the CR group. There was a statistically significant difference in survival distributions for the CR versus control (ad lib) group. *p* < 0.05 by Kaplan-Meier survival analysis.

### CR improves neurological recovery after asphyxial cardiac arrest and resuscitation

To evaluate neurological recovery post-CA of rats that successfully resuscitated and survived, we measured NDS at 4, 24, 48, and 72 h post-ROSC. As shown in Table 1, NDS testing assesses arousal, brainstem reflexes, basic motor strength, withdrawal (sensation), gait, and primitive behaviors. In a pre-experimental, healthy state, rats have perfect NDS scores of 70, whereas post-CA all rats exhibit deficits in NDS.

As shown in Figure 3, the CR group exhibited significantly higher NDS scores in comparison to the control group at every assessed timepoint post-CA. Analysis by two-way repeated measures ANOVA revealed an overall group difference (p<0.05), and significant differences between the control and CR group at 4 h (16.1±5.0 [control n=12] versus 21.7±4.7 [CR n=14]; p<0.05), at 24 h (49.2±9.3 [control n=11] versus 56.9±6.9 [CR n=14]; p<0.05), at 48 h (50.3±12.4 [control n=10] versus 64.2±3.9 [CR n=14]; p<0.05), and at 72 h (52.4±12.5 [control n=10] versus 67.2±2.5 [CR n=14]; p<0.05), by post-hoc t-tests. These results further suggest that a neuroprotective mechanism against ischemic insult appears to manifest as a downstream consequence of 14-hrs of caloric restriction, which in turn augments neurological recovery.

**FIG. 3.**
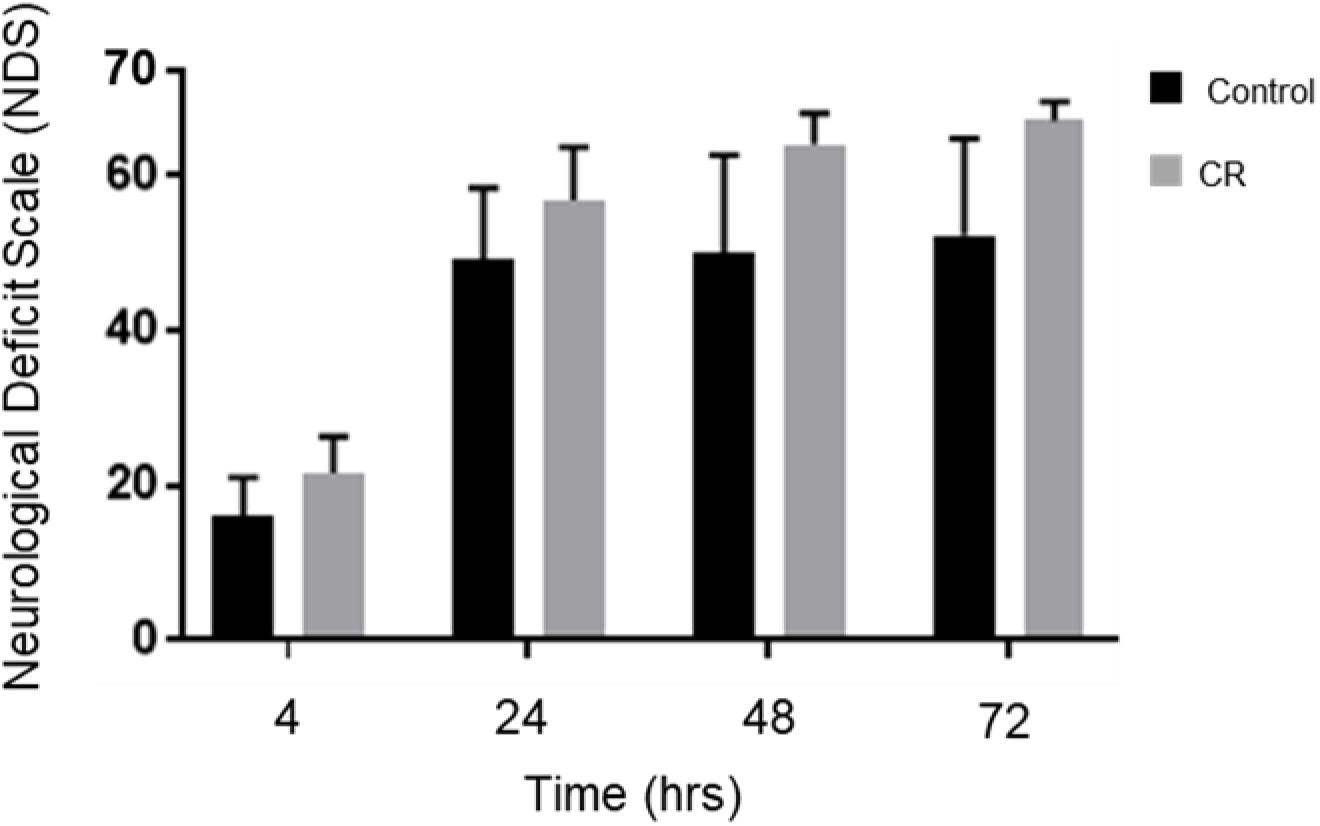
Neurological recovery of post-CA rats. NDS was measured at 4,24,48, and 72 h post-CA. Rats in the CR group consistently scored significantly higher at all measured timepoints. *p<0.05 by two-way analysis of variance with post-hoc t-tests. CA, cardiac arrest; CON, control; NDS, Neurological Deficit Scale; SEM, standard error of the mean.

### CR reduces neurodegeneration in multiple brain regions

To examine the neuroprotective capacity of overnight caloric restriction at the cellular level, we utilized Fluorojade-B (FJ-B) staining to scan and identify variance in neuronal degeneration between the control and CR group at 72h post-CA. Images of the areas with FJ-B positive neurons were captured (as indicated by the red squares in Fig. 4A and the representative images for each region in Fig. 4B). Demonstrative of global cerebral ischemia, several physiological regions throughout the brain were identified to have FJ-B positive neurons. All regions were quantified (as previously described) and three locales were statistically determined to have significant difference between groups. Remarkably, at 72h post-CA, these regions showed zero FJ-B positive neurons in the CR group (n=6). Comparatively, in the control group FJ-B positive neurons were prevalent in a region of the subiculum (7.3±6.7 cells/mm^2^ [n=6]; p<0.05), a region of the principal sensory nucleus (1.7±1.6 cells/mm^2^ [n=6]; p<0.05), and a region of the spinal trigeminal nucleus (1.9±1.6 cells/mm^2^ [n=6]; p<0.05). The subiculum, which is a component of the hippocampal formation, has been linked to processes of spatial navigation and awareness^24^; the principal sensory nucleus plays a role in sensation and proprioceptive feedback from whiskers and the muscles of the face^25^; and the spinal trigeminal nucleus receives sensory information from several cranial nerves^26^. These results suggest that 14 h of caloric restriction prior to ischemic insult leads to complete neuroprotective effects in certain physiological locales of the brain that may play contributory roles in arousal, awareness, and functions that are vital to survival.

**Figure 4.**
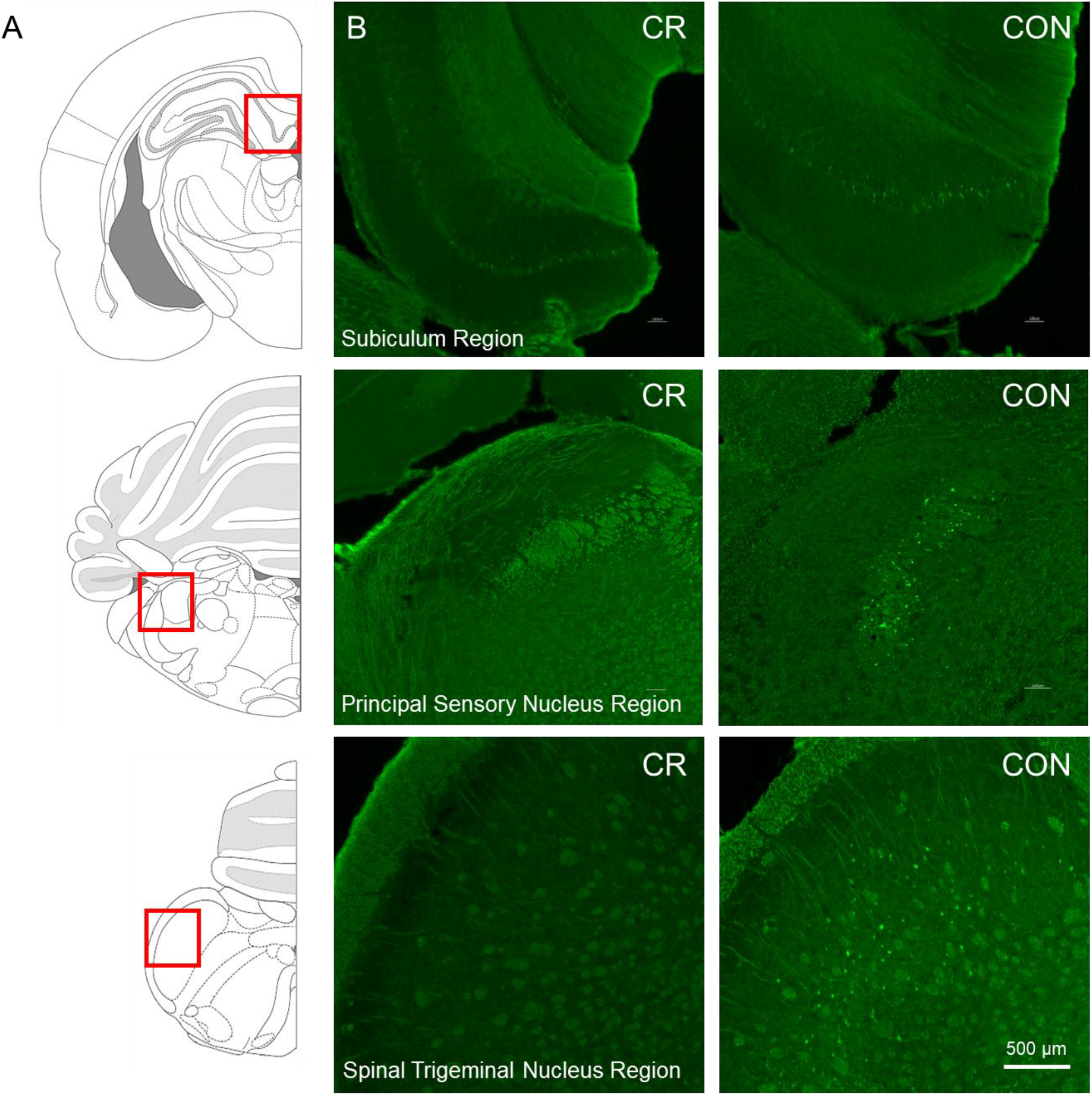
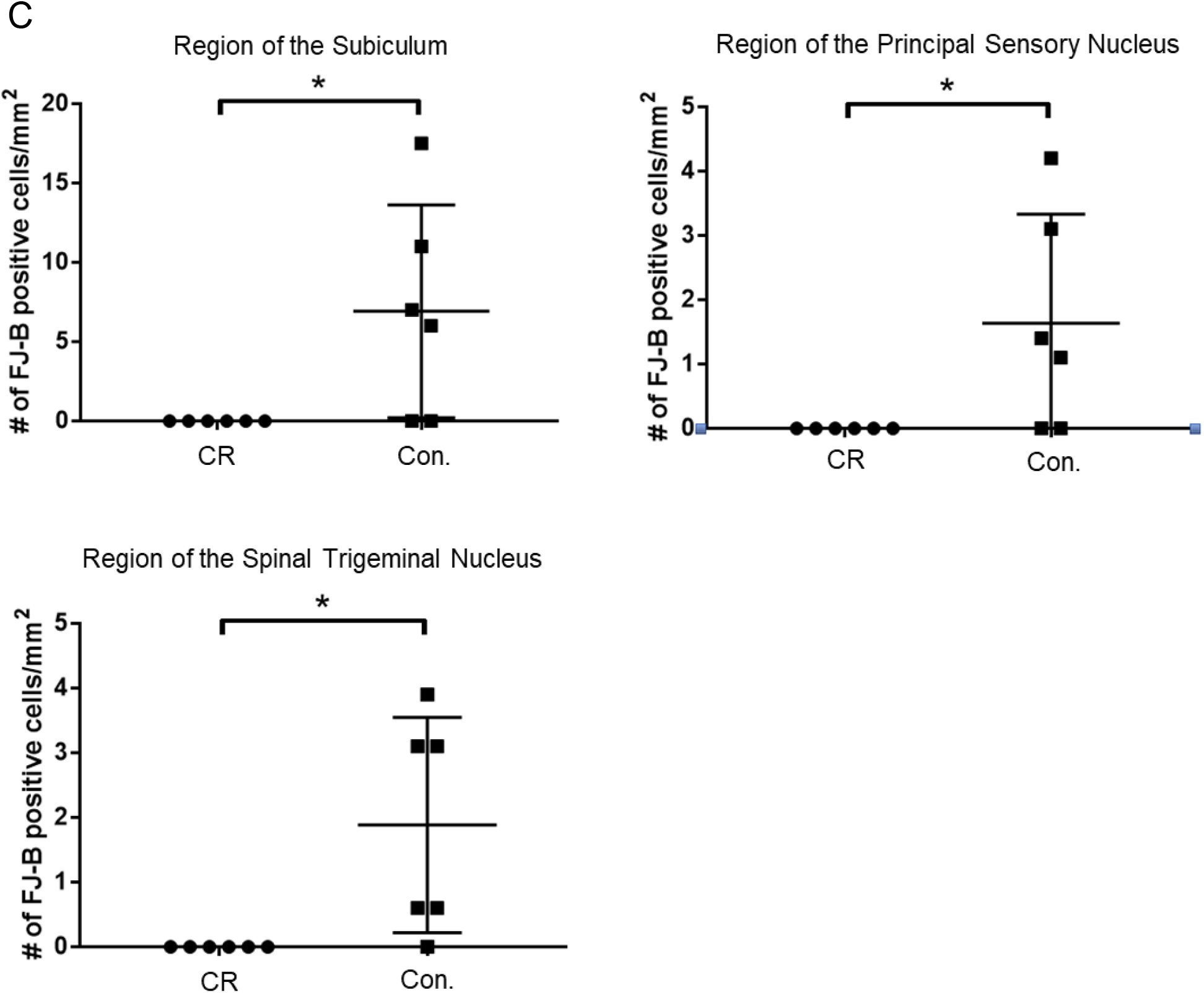
Neurodegeneration in multiple brain regions at 72h post-CA. Neurodegeneration in multiple brain regions at 72h post-CA. (**A**) Brain atlas. The region within the red square displays the physiological locales of neurodegeneration as shown in (B). (**B**) Brain sections were stained with FJ-B; positive neurons are fluorescent (green). Images were cropped and signal intensities were adjusted linearly to be optimal for demonstration. (**C**) Number of FJ-B positive neurons were counted from 3 brain sections per region in each animal. The number of FJ-B positive neurons were significantly higher in control rats; CR rats exhibited no FJ-B positive neurons at these regions. *p<0.05 by unequal variances t-test. CA, cardiac arrest; FJ-B, Fluorojade-B, CON, control; SEM, standard error of the mean.

### CR leads to ketosis

Given that both glycemia and caloric restriction are implicated in ketone body production, we measured capillary blood ketone levels (β-hydroxybutyrate) after 14-hrs of caloric restriction. As shown in Figure 5, the CR group exhibited significantly higher blood ketone levels (mean ± standard deviation [SD], 1.4 ± 0.5 mmol/L; n = 12) in comparison to control (0.6 ± 0.2 mmol/L; n = 12; *p*<0.01). These results indicate that 14-hrs of caloric restriction is of sufficient length to significantly increase the production of endogenous blood ketone levels.

**FIG. 5.**
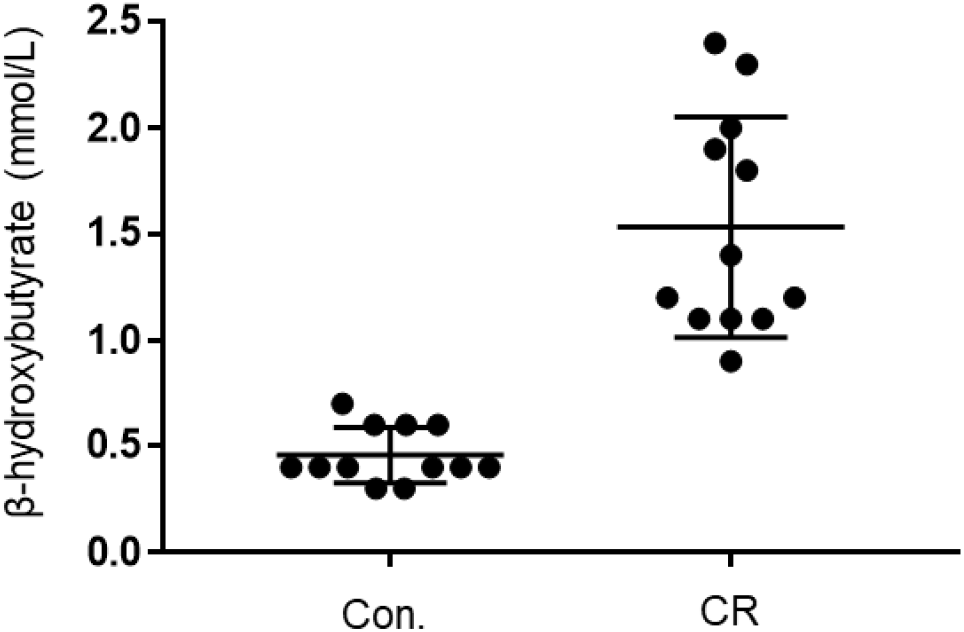
Capillary blood ketone (β-hydroxybutyrate) levels of control vs. CR rats. Blood ketones were measured after 14-hrs of caloric restriction. CR significantly increases ketone levels. ** *P* < 0.01; by unpaired t-test. CR, caloric restriction; CON., control; SEM, standard error of the mean.

### CR leads to higher corticosterone and lower glucagon and insulin

As an indicator of stress, and to further elucidate upon the potential role of glycemia as a contributable outcome factor, we assessed corticosterone, glucagon, and insulin levels in arterial blood collected after 14-hrs of caloric restriction. As shown in Figure 6, the CR group exhibited significantly higher corticosterone levels in comparison to the control group (398,542 ± 59,024 pg/mL [n = 12] versus 196,040 ± 22,092 pg/mL [n = 12]; *p*<0.05), likely indicative of a stress-response following metabolic deficits. Blood levels of glucagon, on the contrary, were significantly lower in the CR group in comparison to the control group (19.6 ± 7.8. pg/mL [n = 9] versus 8.4 ± 2.9 pg/mL [n = 9]; *p*<0.05). Blood levels of insulin were likewise significantly lower in the CR group in comparison to the control group (2621 ± 965.4 pg/mL [n = 8] versus 393.4 ± 265.9 pg/mL [n = 8]; *p*<0.05).

**FIG. 6.**
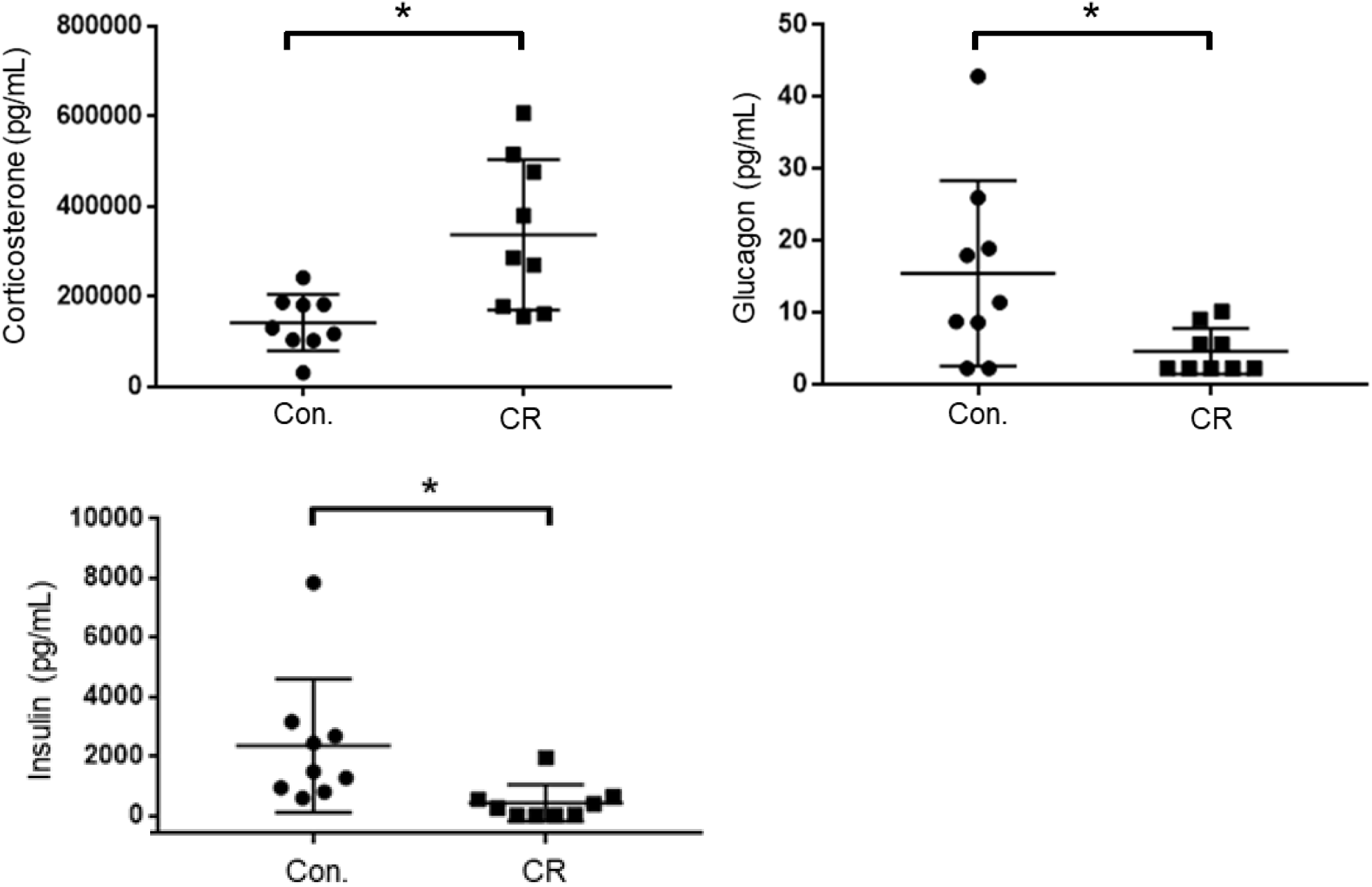
Arterial blood corticosterone, insulin, and glucagon levels of control vs. CR rats measured after 14-hrs of caloric restriction. CR significantly increases corticosterone levels and lowers glucagon and insulin levels. * *P* < 0.05; by Welch’s t-test. CR, caloric restriction; CON, control; SEM, standard error of the mean.

### CR does not change expression of SIRT1 and BDNF

Caloric restriction is known to upregulate brain-derived neurotrophic factor (BDNF) and sirtuin 1 (SIRT1) pathways in the brain, particularly following subacute periods of dietary restriction. To assess the potential upregulation of 14-hrs of caloric restriction, we conducted Western blot analyses on brain homogenates of a separate cohort of rats that were calorically restricted for 14-hrs. Surprisingly, as shown in Figure 7, we found no significant difference in either BDNF or SIRT-1 expression after 14-hrs of caloric restriction in comparison to controls.

**FIG. 6.**
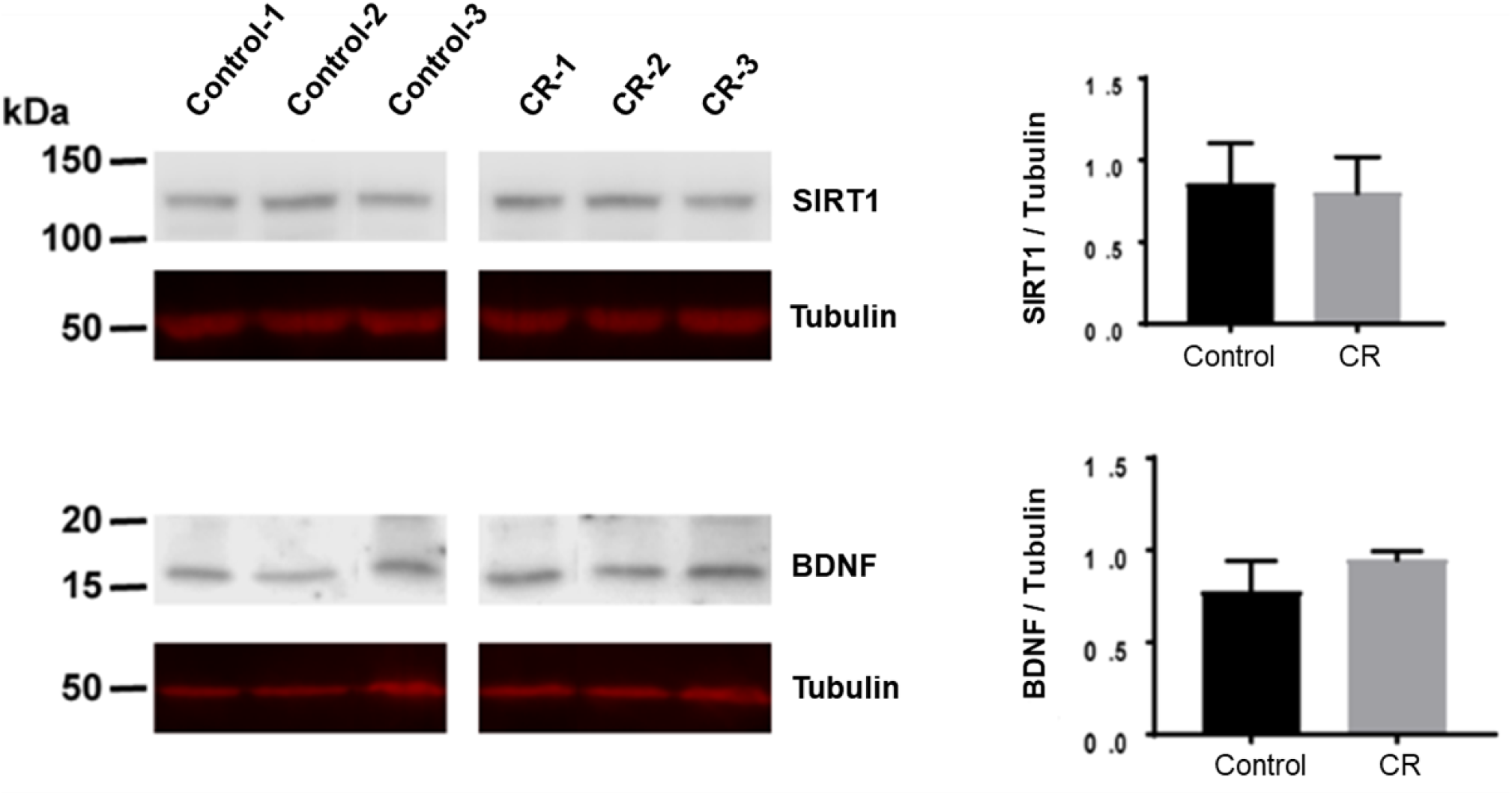
BDNF and SIRT-1 in brain homogenates following 14-hrs of CR. (**A**) The Western blot was probed with SIRT1 antibody (AHP1003) and showed no difference between CR (n=3) and control groups (n=3; p=0.78). It was then stripped and reprobed with an anti-tubulin antibody to confirm loading equivalence. (**B**) The Western blot was probed first with BDNF antibody (AHP1003) and showed no significant difference between CR (n=3) and controls (n=3; p=0.16). It was then stripped and reprobed with an anti-tubulin antibody to confirm loading equivalence. Lane 1-3 are control samples, Lane 3-6 are CR samples.

## Discussion

To date, many studies have indicated that CR paradigms exert robust protection against a multitude of diseases, including ischemic injury. Missing from the literature, however, is whether ultra-short (e.g. overnight) CR prior to a major ischemic insult provides protection. Such an ultra-short CR paradigm provides vastly more translational value, potentially introducing new therapeutic directions, while also having implications for experimental science and reproducibility. In this study, we investigated the effect of a transient 14-hr period of 75% overnight CR on outcome and neurological recovery after asphyxial CA, which leads to a non-shockable form of CA, now the most common type of cardiac arrest^27–30^. Our results reveal improved survival, improved neurological recovery, and potent neuroprotection in specific regions of the rodent brain, including the subiculum, principal sensory nucleus, and spinal trigeminal nucleus. In attempt to dismantle the potential mechanisms involved at large, we explored various candidates that have been linked in prior studies with neuroprotection downstream of short-term caloric restriction. Our findings yield a normalization of blood glucose, increase in β-hydroxybutyrate and corticosterone, decrease in glucagon and insulin, and no significant upregulation of SIRT-1 and BDNF. Although further studies are warranted to pinpoint the specific mechanisms of overnight CR, to our knowledge, this is the first study showing significant benefit of overnight CR in an acute ischemic injury model. Below, we aim to briefly review the potential candidate mechanisms and related caveats in light of overnight CR potentially acting via an assembly of pathways that collectively convene to yield the noted effects of recovery in our model.

### CR-induced normoglycemia

Hyperglycemia has been widely shown to exacerbate various models of cerebral ischemia^31–33^. In humans, it has been associated with significantly higher morbidity and mortality, and reduced long-term recovery^32^. Blood glucose levels have also been shown to significantly rise during CPR, possibly due to a stress-induced mechanism that leads to a deleterious hyperglycemic onset that further worsens neurological outcome^34^.

In our model, our rats exhibit slight stress-induced hyperglycemia after the surgical preparation needed to prepare for the CA experiment. Our findings reveal that a 14-h period of overnight CR is ample to blunt and in fact normalize this surgical stress-induced hyperglycemia prior to cardiac arrest induction. Furthermore, this stress-induced hyperglycemia in control (*ad lib*) rats is worsened after CA+CPR, when glucose levels rise even higher, mirroring the stress-induced hyperglycemic phenomena noted in humans after CPR. Meanwhile, CR rats do not display any form of hyperglycemia during the stress of surgical preparations nor following CPR. It appears that a brief 14-h period of CR is sufficient to completely normalize the hyperglycemic response in our experimental paradigm. We cannot rule out whether the improvement in neurological outcome after CR is a result of normalization of blood glucose levels. However, it has been reported in both human and rodent studies that normalization of hyperglycemia by insulin administration did not improve neurological outcome after traumatic brain injury^35^. We therefore postulate that the mechanisms underlying the neuroprotective effects of CR in our model go beyond those afforded by normoglycemia, warranting further experimental investigation.

### CR-induced ketosis

Global cerebral ischemia leads to a variety of deleterious effects, primarily due to a decrease in oxidative metabolism via a reduction of oxygen availability. Ketone bodies have been widely studied as neuroprotective molecules with promising potential to ameliorate downstream ischemic injury, such as those caused by lactate generation, apoptotic cascade activation, and free radical proliferation^36^. Administration of β-hydroxybutyrate in particular has been shown to prolong the survival time in rodent models of global cerebral ischemia, and to reduce infarct size following focal cerebral ischemia^37^. Moreover, a study examining the effects of a ketogenic diet on CA-induced cerebral ischemia showed that induction of endogenous ketone bodies led to complete neuroprotection in certain locales of the brain^38^.

In our model, 14-h of overnight CR induced a significant rise in endogenous β-hydroxybutyrate levels. It is significant to note that the blood-brain barrier (BBB) is relatively impermeable to ketone bodies unless they are transported by the carrier protein monocarboxylic acid transporters (MCT1 and MCT2). Although it is not well understood how MCTs are regulated, CR has been shown in various studies to increase the BBB uptake of ketone bodies by upregulating expression of MCT1^39^. In addition, studies have demonstrated an increase in MCT1 and MCT2 expression following ischemic cerebral injury^40^. In consideration of these studies collectively, we postulate that the neurological recovery exhibited in our model of CA may, in part, be influenced by a β-hydroxybutyrate-mediated mechanism that warrants further investigation.

### Corticosterone, glucagon, and insulin

Cortisol is the primary glucocorticoid found in humans, whereas corticosterone is the dominant glucocorticoid in rodents. Glucocorticoids have been shown to have bimodal affects that may be beneficial during acute rises but harmful after prolonged elevation^41^. Additionally, they have both pro- and anti inflammatory effects in various brain injury models^42^. They are believed to be potent inducers of apoptosis mainly through a shutdown of inflammatory response via inhibition of the NF-κB, a pro-inflammatory transcription factor^43^. In other studies, they have been linked with deleterious effects that further exacerbate neurological recovery following brain injury^44^. Given some of the deleterious effects of elevations in glucocorticoids while caloric restriction, with its vastly reported benefits in the literature, leads to elevations in glucocorticoids, the “glucocorticoid paradox of caloric restriction” has been proposed^45^. Indeed, knocking down the elevation of glucocorticoids during caloric restriction has been shown to lead to even higher neuroprotection. This suggests that benefits of caloric restriction may not be mediated by elevations of glucocorticoids.

Insulin and glucagon work in tandem to balance blood glucose levels within the body. Likewise, various studies have shown glucagon and insulin to share similar mechanisms of neuroprotection^46^. In a rodent model of ischemic brain injury, insulin and glucagon improved poststroke outcome in animal models by decreasing glutamate in the circulation and cerebrospinal fluid^47^. In our model, 14-hrs of CR led to higher serum corticosterone levels and lower glucagon and insulin levels. We postulate that the increase in corticosterone may be a downstream effect of very short-term dietary-induced stress, which may mediate some form of preconditioning. Moreover, the decrease in glucagon and insulin levels may be a potential reflection of the onset of glycemic normalization in the CR rats, and may loosely suggest that neither are directly involved in the yielded effects of a 14-hr period of CR. The intertwining and role of these metabolites in the grander mechanisms of overnight CR remains nebulous and warrants careful examination.

### SIRT-1 and BDNF

SIRT1 is a NAD+-dependent deacetylase which has a multitude of downstream effects, including inhibition of NF-kB, protection from oxidative stress, and a regulator of autophagy. SIRT1 has also been shown to possess neuroprotective properties in a variety of pathological conditions including neurodegenerative diseases and cerebral ischemia. Moreover, cerebral ischemia injury has been shown to be attenuated by short-term caloric restriction via upregulation of SIRT1 expression^11,16^. Despite these overwhelming studies in lengthier CR experiments, our results yielded no significant difference in the upregulation of SIRT1 following a 14-h period of CR.

BDNF is a neurotrophic factor with a wide variety of immediate and long-term effects. It exerts immediate effects by altering synaptic transmission, while exerting longer term effects via synaptogenesis and neurogenesis^18^. Through these mechanisms, BDNF and its receptor tropomyosin-related kinase B (TrkB) provide neuroprotection against stress and cell death. Long-term CR is known to have a multitude of beneficial effects, including protection against age-related cognitive decline^11^. Increased BDNF expression has been shown to mediate many of these effects^48^. Likewise to our findings with SIRT1, our results yielded no significant difference in BDNF expression following such a transient period of CR. We therefore suggest that the mechanisms involved in such an ultra-short overnight CR are independent of those afforded by SIRT-1 and BDNF, contrary to the supporting literatures that investigate lengthier periods of CR.

### Implications for basic science research

In the clinical setting, overnight fasting is implemented preoperatively in attempt to prevent pulmonary aspiration of digestive content during endotracheal intubation and while under the effects of general anesthesia. This practice has been standardized, as even the most minor cases of patient aspiration could lead to aspiration pneumonitis, pneumonia, and respiratory failure. In many animal models of experimental research, however, preoperative overnight fasting is seldom implemented. In rodent models, justification for this is in part due to a lack of emetic reflex in mice and rats, which diminishes aspiration-related complications^49^.Yet, reports about laboratory practices the night prior to experimentation range from ad lib diet, overnight fasting, to unspecified information in the materials and methods of published studies. In rodent cardiac arrest models specifically, some studies report complete ad lib diet^50–53^, while others report fasting the night prior to experiments^54–56^, and yet others do not specify this information^57–60^. Our findings suggest that such lack of heterogeneity towards certain pre-experimental conditions, in this case an extremely transient dietary adjustment, may lead to incongruity of results and lack of reproducibility within the scientific community. As such, we strongly suggest that careful consideration be made towards the justification and proper reporting of pre-experimental dietary methodologies.

### Limitations

First, although we investigated key candidate markers in select pathways shown to be important in CR and various models of cerebral ischemia, our study does not examine a multitude of other potential pathways and metabolites involved in the mechanisms of ultra-short caloric restriction. We therefore suggest that future transcriptomic and metabolomic studies may be of great benefit towards untangling the specific mechanisms involved herein. Experimental manipulations must be done to tease apart the mechanisms underlying such drastic effects exhibited by a brief fasted state. Additionally, further support for our findings would require additional histological investigation (beyond FJ-B staining) with formal stereological analysis, additional functional assays and an expansion of sample size. Lastly, because rodents are nocturnal creatures, we are not certain whether overnight fasting in rodents may be equivalent to daytime fasting in diurnal creatures, including humans. Rodents also have significantly higher metabolic rates than larger animals. Certainly, studies have shown that overnight CR of rodents is incomparable to overnight CR in humans^61^. Given that many physiological parameters are regulated by circadian rhythm and metabolic rate, the onset and amount of CR at various points in the circadian rhythm may have disparate implications for different species.

## Conclusion

To our knowledge, this is the first study focusing on ultra-short CR (< 24h) in an ischemic brain injury model. We demonstrate that a mere 14-hr period of overnight caloric restriction prior to cardiac arrest and resuscitation in a rodent model has significant effects on survivability and neurological recovery, as supported by behavioral assessments in addition to histological and blood serum analyses. Remarkably, our histological assessments show up to complete neuroprotection in certain locales of the brain. These findings motivate future studies to pinpoint the specific mechanisms of such a potent ultra-transient caloric restriction and spotlights the critical necessity of standardizing pre-experimental dietary methodologies to ensure research homogeneity and reproducibility.

## Acknowledgments

Special thanks to Suhas Sureshchandra and Dr. Ilhem Messaoudi-Powers, University of California (UC) at Irvine, for their support with the Luminex MAGPIX system; to Francisco Aguirre, Ramin Badiyan, Natalie Khalili, Jeannine C. Wang, Tin Dinh, Juan Alcocer, Stephanie Kim, and Omid Mirkhani, for experimental assistance; and to Michael Klymkowsky, who developed the monoclonal antibody E7 directed against β-tubulin, which was obtained from the Developmental Studies Hybridoma Bank developed under the auspices of the National Institute of Child Health and Human Development and maintained by the Department of Biological Sciences, University of Iowa, Iowa City, IA.

This work was supported by grants awarded to Dr. Yama Akbari, including: NIH R21EB024793, 5KL2TR0001416, a Faculty Research Grant from the UC Irvine School of Medicine Committee on Research and Graduate Academic Programs, as well as funds from the UC Irvine School of Medicine and Department of Neurology. Additionally this work was supported by the Roneet Carmell Memorial Endowment Fund.

