## Supplemental Figure 1 for "Overnight caloric restriction prior to cardiac arrest and resuscitation leads to improved survival and neurological outcome in a rodent model"

Supplemental Figures

| **Arterial Blood Measurement** | **Control** ***Pre-CA*** | **CR** ***Pre-CA*** | **Control** ***Post-CA*** | **CR** ***Post-CA*** | **Significance** |
|---|---|---|---|---|---|
| pH | 7.42 ± 0.06 | 7.39 ± 0.05 | 7.40 ± 0.04 | 7.43 ± 0.09 | None, P>0.05 |
| $CO_2$(mmHg) | 38.3 ± 7.66 | 39.1 ± 7.46 | 38.9 ± 7.64 | 39.2 ± 6.11 | None, P>0.05 |
| $O_2$(mmHg) | 166.4 ± 34.5 | 165.42 ± 24.5 | 164.9 ± 32.5 | 169.5 ± 55.7 | None, P>0.05 |
| $HCO_3^-$(mmol/L) | 24.9 ± 2.2 | 25.9 ± 2.2 | 25.9 ± 2.1 | 26.0 ± 2.2 | None, P>0.05 |
| Sodium (mmol/L) | 139.7 ± 2.3 | 140.1 ± 3.0 | 139.1 ± 2.5 | 139.2 ± 1.2 | None, P>0.05 |
| Potassium(mmol/L) | 3.7 ± 0.4 | 3.8 ± 0.5 | 3.6 ± 0.6 | 4.0 ± 0.3 | None, P>0.05 |
| Calcium(mmol/L) | 1.37 ± 0.06 | 1.34 ± 0.07 | 1.37 ± 0.08 | 1.38 ± 0.06 | None, P>0.05 |
| Hemoglobin (g/dL) | 11.8 ± 0.9 | 11.5 ± 0.8 | 11.9 ± 0.7 | 12.0 ± 1.2 | None, P>0.05 |

**Figure S1.** Arterial blood was collected and analyzed in both control and CR groups 10 minutes prior to CA and after CA. No association was found between groups in pre- or post-CA arterial blood gas measurements (e.g. pH, CO2, O2, electrolytes, etc.).
